# Measuring cystic fibrosis drug responses in organoids derived from 2D differentiated nasal epithelia

**DOI:** 10.1101/2021.07.20.453105

**Authors:** Gimano D. Amatngalim, Lisa W. Rodenburg, Bente L. Aalbers, Henriette H. M. Raeven, Ellen M. Aarts, Iris A.L. Silva, Wilco Nijenhuis, Sacha Vrendenbarg, Evelien Kruisselbrink, Jesse E. Brunsveld, Cornelis M. van Drunen, Sabine Michel, Karin M. de Winter-de Groot, Harry G. Heijerman, Lukas C. Kapitein, Magarida D. Amaral, Cornelis K. van der Ent, Jeffrey M. Beekman

## Abstract

Cystic Fibrosis (CF) is caused by genetic defects that impair the cystic fibrosis transmembrane conductance regulator (CFTR) channel in airway epithelial cells. These defects may be overcome by specific CFTR modulating drugs, for which the efficacy can be predicted in a personalized manner using 3D nasal-brushing-derived airway organoids in a forskolin-induced swelling assay. Despite of this, previously described application of 3D airway organoids in CFTR function assays have not been fully optimal. In this report we therefore describe an alternative method of culturing nasal brushing-derived airway organoids, which are created from an equally differentiated airway epithelial monolayer of a 2D air-liquid interface culture. In addition, we have defined organoid culture conditions, with the growth factor/cytokine combination neuregulin-1β and interleukine-1β, which enabled consistent detection of CFTR modulator responses in nasal airway organoids cultures from subjects with CF.

## Introduction

Cystic fibrosis (CF) is a monogenic epithelial disease caused by mutations in the cystic fibrosis conductance transmembrane regulator (*CFTR*) gene ^1^. This defect impairs CFTR-dependent anion conductance in airway epithelia ^2^, which leads to a severe respiratory disease ^3^. CFTR modulators are target-specific drugs that may restore CFTR function in individuals with CF ^4^. However, the efficiency of modulators largely depends on the CFTR genotype of an individual with CF. More than 2,000 distinct CFTR mutations have been reported (http://www.genet.sickkids.on.ca//) with variable effects on CFTR expression or function. In addition to common mutations, such as the F508del allele, approximately 1,000 rare mutations have been identified that each affect less than 5 individuals worldwide. This low prevalence makes it unfeasible to determine CFTR modulator drug efficacy in large cohort clinical studies.

As an alternative of determining drug efficacy directly in individuals with CF, the effects of CFTR modulators can be predicted using patient-derived epithelial cultures in functional CFTR assays ^5^. This is traditionally done with 2D air-liquid interface (ALI)-differentiated airway epithelia by assessment of CFTR-dependent chloride (Cl^-^) conductance via electrophysiology ^6–8^. However, a major disadvantage of the ALI-culture model system is the limited scalability. In contrast, CFTR-expressing epithelial organoids from various tissues, i.e. airway, intestine, kidney, are emerging as a novel model system in which drug efficacy can be tested more efficiently in a mid-to high-throughput fashion ^9–11^. Previously, we and others have shown that intestinal organoids from subjects with CF can be used to predict drug responses in a forskolin-induced swelling (FIS) assay, reflecting CFTR-dependent fluid secretion ^12–15^. Nevertheless, based on the origin of CF respiratory disease, it remains postulated that airway epithelial models are more predictive for determining CFTR modulator responses.

Indeed, we previously reported CFTR modulator response measurements in long-term expanded distal airway organoids using FIS ^11^. However, we observed large variations in FIS measurements, as swelling was limited to well-differentiated spherical structures. Others have successfully used 3D nasal airway organoids (NAOs) derived from minimal-invasive nasal brushings in functional CFTR assays ^16–18^, which are a more suitable option for personalized drug testing compared to cultures derived from invasive (tracheo)bronchial and intestinal tissues. However, previously described functional CFTR assays using NAOs were either low-in throughput or required CFTR function measurements over large time periods ^16–19^. Altogether, there is a remaining need for a further optimized and scalable FIS assay using airway organoids, especially derived from nasal brushings, which enables efficient CFTR modulator response measurements in subjects with CF. In this report, we describe an alternative organoid culture method, in which NAOs are derived from 2D differentiated human nasal epithelial cell (HNEC) monolayers. We furthermore describe optimized airway organoid culture condition to improve CFTR modulator response measurements, by including the growth factor/cytokine combination neuregulin-1β and IL-1β. Validation studies using this culture condition showed consistent detection of genotype-specific responses to CFTR modulators in FIS assays, including repairing effects of the FDA-approved CFTR triple modulator therapy VX-661/VX-445/VX-770 ^20^.

## Results and Discussion

We previously reported CFTR function measurements in distal airway organoids using a FIS assay ^11^. However, in this study we observed large variation in swelling between individual organoids, which was restricted to large spherical structures. This is likely caused by the unsynchronized differentiation of individual organoids, which may influence differentiation-dependent CFTR function (Figure S1). To overcome this unsynchronized differentiation of organoids within a culture, we set up a novel approach in which evenly differentiated airway organoids are established from a 2D differentiated human nasal epithelial cells (HNEC) monolayer (Figure 1A). In the organoid culture procedure, HNEC derived from nasal brushings are first isolated and expanded in regular 2D cell cultures (Figure 1B). HNEC stained positive for p63 and cytokeratin 5 (KRT5), confirming a basal stem cell phenotype (Figure 1C). After expansion, HNEC were differentiation in conventional 2D air-liquid interface (ALI) transwell-cultures, to recapitulate the mucociliary airway epithelium. Indeed, well-differentiated ALI-HNEC cultures resembled the native nasal epithelial tissue, by displaying a similar pseudostratified morphology and consisting of MUC5AC/CC10^+^ secretory cell, β-tubulin IV^+^ ciliated cells, and p63/KRT5^+^ basal cells (Figure 1D and E). CFTR function and effects of modulators were confirmed in ALI-HNEC from a healthy control (HC) and individual with CF and a F508del/F508del genotype (Figure S2). During isolation of airway organoids from resected airway tissues, based on the method described by Sachs et al.^11^, we observed that large epithelial fragments, obtained after collagenase treatment, self-organized into well differentiated organoids within a few days after gel embedding. Based on this observation and the assumption that ALI-cultures reflect the differentiated airway epithelia tissue, we proposed that epithelial fragments from 2D-cultures could be converted into 3D organoids as well. Indeed, embedding of differentiated ALI-derived epithelial fragments in a 3D extracellular matrix led to formation of organoids with clear lumens within 48 h (Figure 1F). From a single 12 mm^2^ transwell insert we are able to generate a yield of organoids that is sufficient for 48 independent wells (approx. 25-50 organoids/well) of a 96 well plate. In terms of scalability, this demonstrates a major advantage of our method compared to the conventional use of 2D ALI-cultures. ALI-culture-derived airway organoids directly displayed a differentiated phenotype. This was confirmed by visual observation of beating cilia and accumulation of mucus (Video S1), as well as by immunofluorescence imaging, demonstrating β-tubulin IV^+^ cilia and MUC5AC^+^ secretory cells inside of the organoids, and p63^+^/KRT5^+^ basal cells at the basal side (Figure 1G, H). Next, we determined whether ALI-derived NAOs could be used to measure CFTR function in FIS assays in a 96 well plate format (Figure 2A). Forskolin stimulation of organoids from HC subjects increased swelling in time, which was dose-dependent and reached a plateau at 5 µM forskolin (Figure 2B-D). Organoid swelling could be attenuated with the NKKC1 inhibitor bumetanide, demonstrating chloride-dependence of forskolin-induced organoid fluid secretion (Figure S3A and B). Moreover, chemical CFTR inhibitors significantly reduced FIS in HC NAOs (Figure 2E and F), indicating CFTR dependence. Upon comparison of cultures from HC subjects and subject with CF, we observed that both HC and CF NAOs displayed cystic lumens, which were not significant different in size (Figure 2G, H). This corresponds with observations made in distal airway organoids ^11^, and suggests intrinsic CFTR-independent fluid secretion mediated by alternative chloride channels. In support of this, stimulation with the Ca^2+^-activated Cl^-^ channel (CaCC) activator E_act_ induced organoid swelling in CF NAOs, which was significant higher compared to HC cultures (Figure S3C and D). This suggests dominance of CFTR-independent fluid secretion in CF NAOs, also observed by others ^21^. In contrast, FIS measured in NAOs from HC subjects was significantly higher compared to subjects with CF (Figure 2I and J), reflecting differences in CFTR-dependent fluid secretion. Although CF and HC NAOs could be distinguished phenotypically based on FIS, we initially did not observe repairing effects of the CFTR corrector and potentiator combination VX-809/VX-770 in CF F508del/F508del NAOs from multiple subjects (Figure S3E and F). Therefore, we aimed to optimize the CFTR modulator response measurements, by modifying the organoid culture conditions. Recent studies suggested that secretory cells are the primary airway epithelial cell type mediating CFTR function ^22,23^. Therefore, we examining a panel of growth factors and cytokines that could modulate secretory cell functions, which were added after plating of the epithelial fragments and during organoid culturing (Figure 3A). The examined factors included, neuregulin-1β (NR), which has been reported to enhance epithelial polarization and differentiation of secretory cells in ALI-cultures ^24,25^. Moreover, the effect of the pro-inflammatory cytokine IL-1β was examined, which has been shown to enhance goblet cell differentiation, CFTR mRNA expression, chloride conductance, and CFTR modulator responses in ALI-cultures ^26–30^. In contrast to independent factors, a combination of NR/IL-1β led in detection of VX-809/VX-770 modulator responses in CF F508del/F508del NAOs (Figure 3B and Figure S4A). NR/IL-1β-cultured HC NAOs displayed a similar swelling response compared to control culture conditions (Figure 3C and S4B). In support of reducing CFTR-independent fluid secretion, NR/IL-1β attenuated lumen formation in CF NAOs, as quantitated by organoids size measurements (Figure S4C and D). Moreover, in line with enhanced specificity of CFTR function, NR/IL-1β increased and reducing the expression of CFTR and the CaCC ANO1 respectively (Figure S4E). The Cl^-^ channel SLC26A9 was also increased upon stimulation with NR/IL-1β, which may support CFTR function, as proposed by others ^31^. NR/IL-1β did not affect the expression of mucociliary differentiation markers (Figure S4F and G).

**Figure 1).**
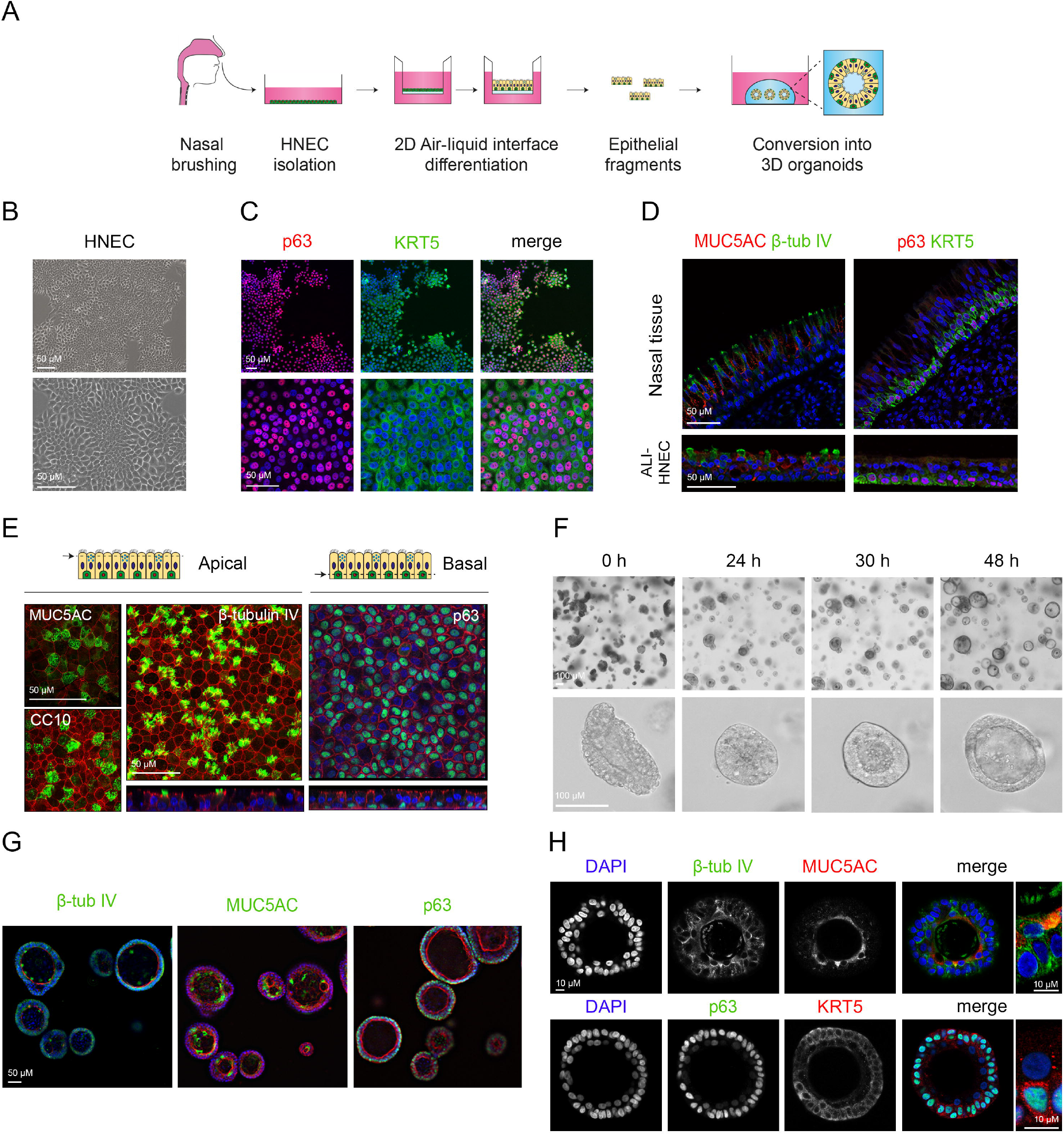
Conversion of differentiated ALI-HNEC cultures into nasal airway organoids. (A) Graphic illustration showing workflow of culturing nasal airway organoids (NAOs) from 2D expanded and differentiated human nasal epithelial cells (HNEC). (B) Brightfield images of a 2D expanded HNEC. (C) IF staining of HNEC with basal cell markers p63 (red) and cytokeratin 5 (KRT5, green). (D) Sections of nasal tissue (top) and 18 days differentiated air-liquid interface (ALI-) cultured HNEC (bottom), demonstrating IF staining of MUC5AC (goblet cells, red), β-tubulin IV (ciliated cells, green), p63 and KRT5 (basal cell markers, green and red respectively). (E) Whole-mount IF confocal images of ALI-HNEC, showing maximal projections of the apical side of the secretory cell markers MUC5AC and Club Cell protein 10 (CC10), and ciliated cell marker β-tubulin IV (Z-stack at the bottom). The projection of the basal side shows the basal cell marker p63 (Z-stack at the bottom). All markers are shown in green. (F) Time course showing self-organization of differentiated ALI-HNEC-derived epithelial fragments into organoids. (G) IF staining of NAOs for β-tubulin IV, MUC5AC, and p63 (all in green). (H) Confocal images showing in the top panels staining with DAPI (blue), β-tubulin IV (green), and MUC5AC (red). The bottom panels show staining with DAPI (blue), p63 (green), and KRT5 (red). Data information: DAPI (blue) was used as nuclear staining (C-E, G, and H). Phalloidin (red) was used as actin cytoskeleton staining (E and G).

**Figure 2).**
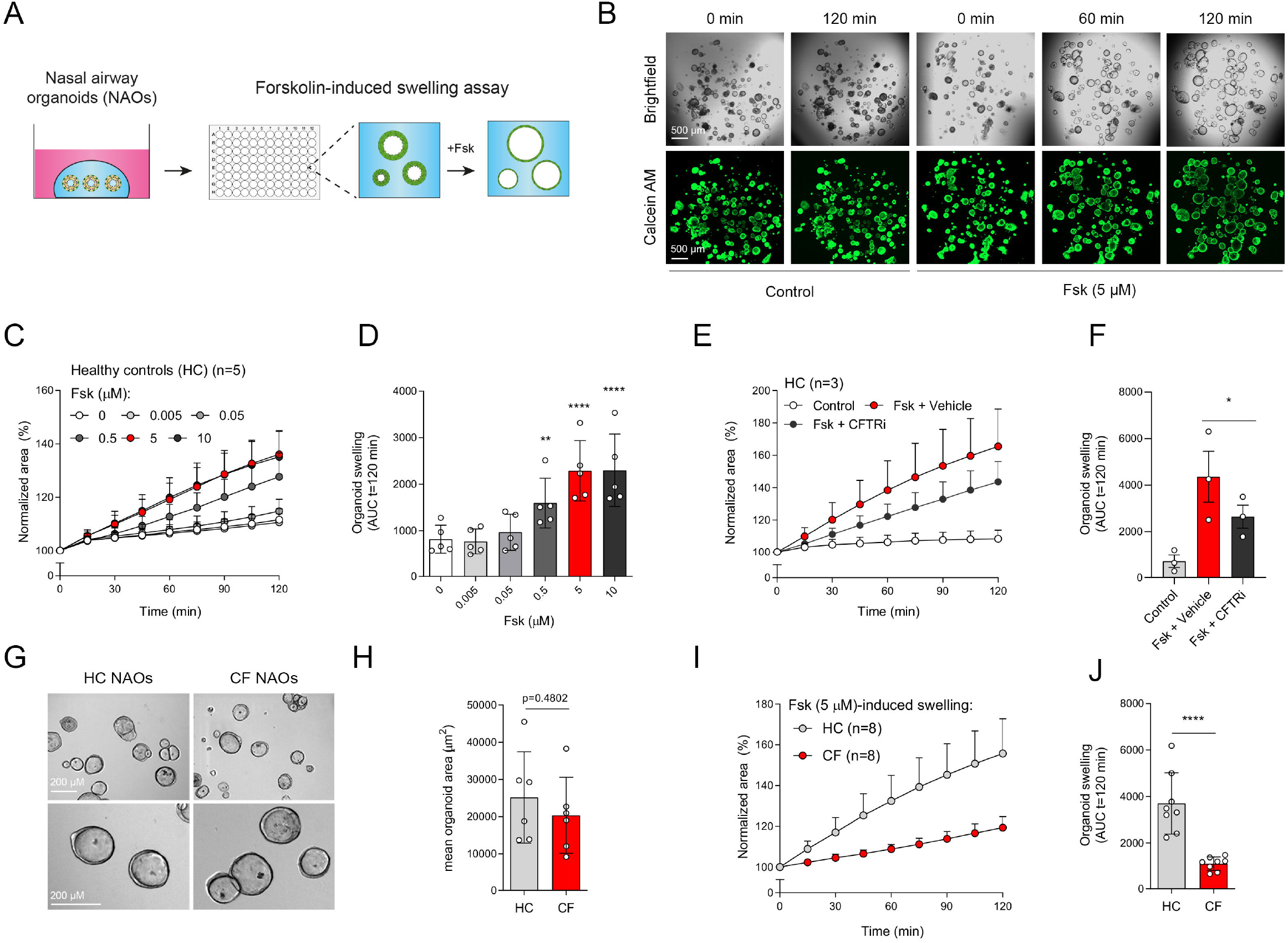
Forskolin-induced swelling assay with ALI-culture derived nasal airway organoids. (A) Graphic illustration showing FIS assay with ALI-derived nasal airway organoids (NAOs). (B) Representative brightfield images (top) and images of calcein green AM esters-strained (bottom) organoids from a HC subject, which were unstimulated (Control) or treated with forskolin (Fsk, 5 µM). Images were taken at t=0, 60, and 120 min after stimulation. (C and D) NAOs from HC subjects (n=5 independent donors) were stimulated with different concentrations of Fsk (0-10 µM), followed by quantification of FIS. (E and F) HC NAOs (n=3 independent donors) were pre-treated with CFTR inhibitors or vehicle, followed by assessment of FIS. (G) Representative brightfield images of NAOs from HC subjects and subjects with CF. (H) Quantification of the mean organoid area (µm^2^, means ± SD) of HC and CF NAOs (both n=6 independent donors). (I and J) Comparison of FIS between HC and CF NAOs (both n=8 independent donors) after Fsk stimulation. Data information: Results are depicted as the percentage change in surface area relative to t=0 (normalized area) measured at 15-minute time intervals for 2 h (means ± SD) (C, E, I), and area under the curve (AUC) plots (t=120 min, means ± SD) (D, F, J). Analysis of differences was determined with a one-way ANOVA and Bonferroni *post-hoc* test (D), and unpaired t-test (F, H, J). ** p<0.01, *** p<0.001, **** p< 0.0001.

**Figure 3).**
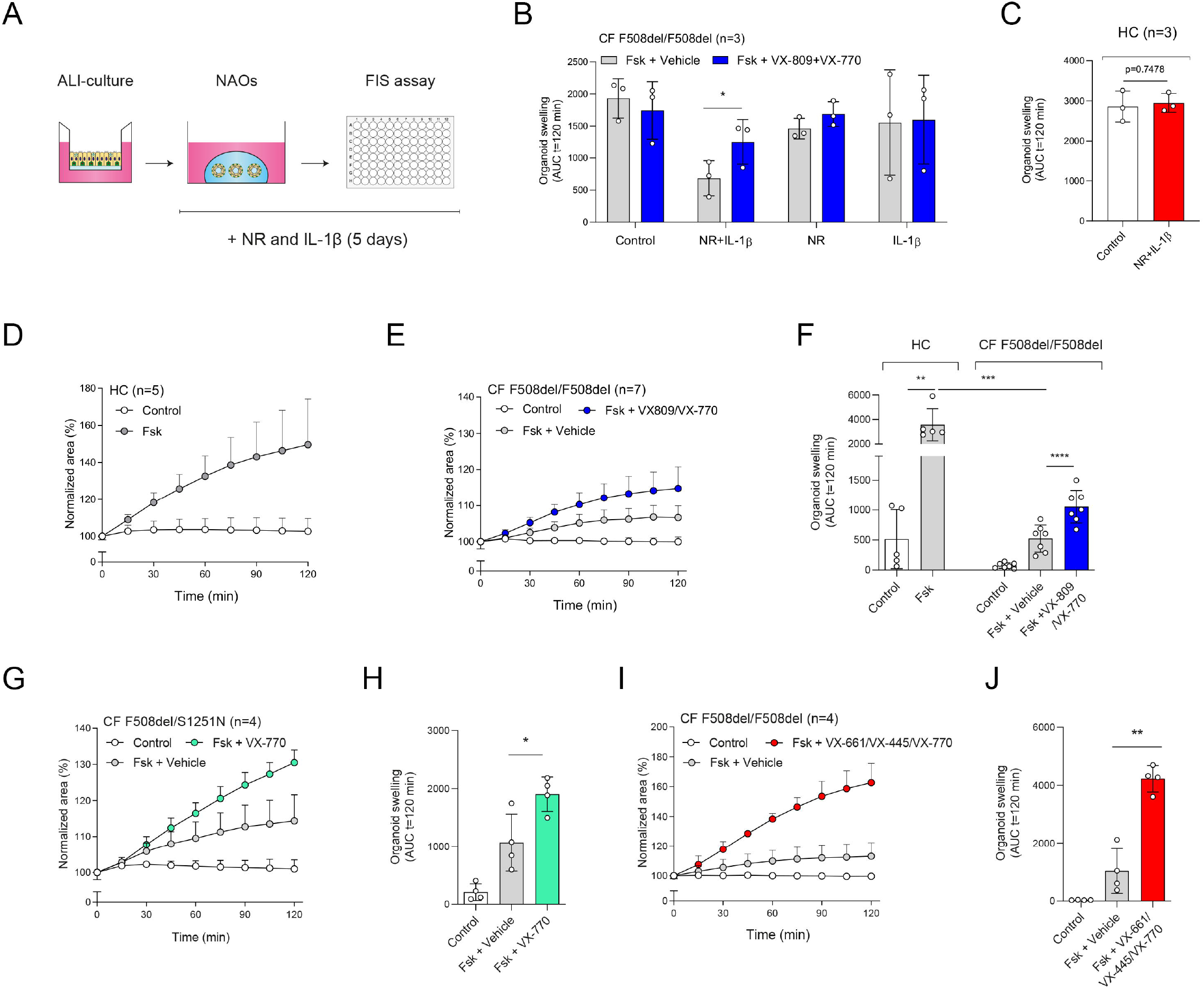
Validation of CFTR modulator responses in NR/IL-1β-cultured NAOs. (A) Illustration showing culturing of NAOs with neuregulin 1β (NR) and interleukin-1β (IL-1β) (B) CF F508del/F508del NAOs (n=3 independent donors) were cultured at control conditions, or with NR, IL-1β, or combination (NR+IL-1β). Afterwards, FIS was determined in response to VX-809/VX-770. (C) Assessment of FIS in HC NAOs (n=3 independent donors) cultured under control conditions or NR+IL-1β. (D) FIS measured in HC NAOs (n=5 independent donors) and (E) CF F508del/F508del NAOs (n=7 independent donors) cultured with NR/IL-1β. FIS responses in CF NAOs were determined in response to VX-809/VX-770. (F) Comparison of FIS measured in HC (Figure 3F) and CF F508del/F508del NAOs (Figure 3E). (G and H) FIS in NR+IL-1β cultured NAOs from CF F508del/S1251N subjects (n=4 independent subjects), stimulated with Fsk and VX-770 or vehicle. (I and J) CF F508del homozygous NAOs cultured with NR/IL-1β (n=4 independent donors) were pre-treated with vehicle or VX-661/VX-445. Swelling was determined afterwards following acute stimulation with Fsk together with VX-770 or vehicle. Data information: Swelling results are depicted as (D, E, G, I) the percentage change in surface area relative to t=0 (normalized area) measured at 15-minute time intervals for 2 h (means ± SD), and (B, C, F, H, J) area under the curve (AUC) plots (t=120 min, means ± SD). Analysis of differences was determined with a paired t-test (B, C, F; within groups, H, J), unpaired t-test (F; HC compared to CF). * p<0.05, ** p<0.01, *** p<0.001, **** p< 0.0001.

We further validated CFTR function and modulator response measurements in the NR/IL-1β culture condition. HC NAOs displayed a significant higher swelling response when compared to cultures from CF F508del/F508del subjects (Figure 3D, F, and S5A). Moreover, the CFTR modulator combination VX-809/VX-770 consistently enhanced FIS measured in CF F508del/F508del NAOs from 7 independent subjects (Figure 3E, F, and S5B). In addition to VX-809, FIS responses in CF F508del/F508del NAOs were modulated upon treatment with other CFTR correctors (Figure S5C). We were also able to detect VX-770 potentiator responses in NAOs from subjects with CF and a S1251N gating mutation (Figure 3G, H, and S5D). Moreover, the highly effective CFTR triple modulator therapy VX-445/VX-661/VX-770 ^20^, induced a high increase in FIS measurements in CF F508del/F508del NAOs cultured with NR/IL-1β (Figure 3I, J, and S5E). VX-445 by itself did not increase FIS, whereas chemical CFTR inhibition completely diminished increases in swelling by VX-445/VX-661/VX-770, demonstrating CFTR-dependence of the modulator treatment effect (Figure S5F and G).

In summary, we described a new method of culturing nasal brushing-derived airway organoids.

Previously we have shown that long-term expanded airway organoids can be cultured as 2D differentiated ALI-cultures ^11^. In this report, we demonstrate further flexibility between 2D and 3D airway culture models, by showing the possibility to convert 2D differentiated ALI-cultures into 3D airway organoids. We furthermore described organoid culture conditions that improved quantification of CFTR modulating drug responses in FIS assays. The NR/IL-1β culture condition reflects the chronically inflamed airway epithelium of individuals with CF, and therefore may act as physiological conditions for testing CFTR modulator responses in NAOs. Further research is required to determine whether NR/IL-1β also improve CFTR modulator responses in other airway model systems, such as distal airway organoids, or other CFTR-expressing epithelial cells. In both the initially tested organoid culture conditions and the optimized conditions with NR/IL-1β, we could not discriminate HC from CF NAOs based on lumen size, which is in contrast to the CFTR-dependent intestinal organoid model ^9,13^. Furthermore, due to limited numbers of donors with diverse CFTR genotypes, we were unable to determine whether residual FIS correlated to disease severity, as shown by others ^18^. In recent studies, other have shown that CFTR modulator responses in 3D nasal cultures correlate with drug responsiveness in individuals with CF ^19,32^. Whether CFTR drug responses measurements in our model correlate with clinical outcome measurements will be determined in follow up studies. CFTR function measurements in ALI-culture derived NAOs may also be used to examine novel target-specific therapies for subjects with CF with unmet, such as assessment of CRISPR-gene editing, read-through agents, or compounds targeting nonsense mediated decay. Moreover, complementary to the widely used 2D ALI-cultures, airway organoids derived from this model may be further used to study other respiratory tract disorders.

## Materials and Methods

### Reagents and Tools Table

| Reagent or Resource | Source | Identifier |
| --- | --- | --- |
| **Antibodies** |  |  |
| Mouse IgG1 anti-β-tubulin IV | Emergo Biogenex | #MU178-UC |
| Rabbit anti-β-tubulin IV | Abcam | #ab179509 |
| Rabbit anti-MUC5AC | Abcam | #ab198294 |
| Rabbit anti-P63 | Abcam | #ab124762 |
| Mouse IgG1 anti cytokeratin 5 | Abcam | #ab17130 |
| Mouse IgG1 anti-CC10 | Acris, Origene | #AM26360PU-N |
| Goat anti-Mouse IgG1, Alexa Fluor 488 | Invitrogen | #A-21121 |
| Goat anti-Mouse IgG1, Alexa Fluor | Invitrogen | #A-21240 |
| 647 |  |  |
| Goat anti-Rabbit IgG, Alexa Fluor 488 | Invitrogen | #A-11034 |
| Goat anti-Rabbit IgG, Alexa Fluor 546 | Invitrogen | #A-11035 |
| **Biological Samples** |  |  |
| Nasal brushings | UMC Utrecht | Protocol ID: 16/586, NL54885.041.16 |
| Nasal turbinate tissue | Academic Medical Center Amsterdam | N/A |
| **Chemicals, Peptides, and Recombinant Proteins** |  |  |
| TrypLE express enzyme | Fisher scientific | # 12605010 |
| Sputolysin | Calbiochem | #560000-10 |
| Collagen IV | Sigma-Aldrich | #C7521 |
| CryoStor® CS10 | StemCell Technologies | #07930 |
| PureCol | Advanced BioMatrix | #5005 |
| Bronchial epithelial cell medium-basal (BEpiCM-b) | Sciencell | #3211 |
| Advanced DMEM F12 | Thermo Fisher | #12634-028 |
| B-27 Supplement, serum free | Thermo Fisher | #12587010 |
| GlutaMAX Supplement | Thermo Fisher | #35050-061 |
| HEPES | Thermo Fisher | #15630080 |
| (±)-Epinephrine hydrochloride | Sigma-Aldrich | #E4642 |
| Hydrocortisone | Sigma-Aldrich | #H0888 |
| 3,3',5-Triiodo-L-thyronine sodium salt | Sigma-Aldrich | #T6397 |
| N-Acetyl-L-cysteine | Sigma-Aldrich | #A9165 |
| Nicotinamide | Sigma-Aldrich | #N0636 |
| SB02190 | Sigma-Aldrich | #S7067 |
| DMH-1 | Selleck Chemicals | #S7146 |
| A83-01 | Tocris | #2939/10 |
| Y-27632 | Selleck Chemicals | #S1049 |
| DAPT | Fisher Scientific | #15467109 |
| TTNPB | Cayman | #16144-1 |
| Recombinant human FGF-7 | Peprtech | #100-19 |
| Recombinant human FGF-10 | Peprtech | #100-26 |
| Recombinant human EGF | Peprtech | #AF-100-15 |
| Recombinant human HGF | Peprtech | #100-39H |
| Recombinant Neuregulin-1β | Peprtech | #100-03 |
| Recombinant Interleukin 1β | Peprtech | #200-01B |
| Penicillin-Streptomycin | Thermo Fisher | #15070-063 |
| Primocin | Invivogen | #ant-pm-2 |
| Amphotericin B | Thermo Fisher | #15290018 |
| Gentamicin | Sigma-Aldrich | #G1397 |
| Vancomycin | Sigma-Aldrich | #SBR00001 |
| Collagenase type II | Thermo Fisher | #17101-015 |
| CFTRinh-172 | Sigma-Aldrich | #C2992 |
| GlyH101 | Sigma-Aldrich | #219671 |
| VX-809 | Selleck Chemicals | # S1565 |
| VX-661 | Selleck Chemicals | #S7059 |
| VX-770 | Selleck Chemicals | #S1144 |
| Correctors C1-18 | Cystic Fibrosis Foundation Therapeutics | https://www.cff.org/Research/Researcher-Resources/Tools-and-Resources/CFTR-Chemical-Compound-Program/ |
| Forskolin | Sigma-Aldrich | #F3917 |
| 3-Isobutyl-1-methylxanthine | Sigma-Aldrich | # I5879 |
| Amiloride | Sigma-Aldrich | #1019701 |
| Calcein green acetoxymethyl (AM) | Invitrogen | #C34852 |
| Alexa Fluor 555 Phalloidin | Invitrogen | #A34055 |
| Phalloidin-iFluor | Abcam | #ab176752 |
| 405 |  |  |
| Prolong Gold antifade reagent | Thermo Fischer | #P36934 |
| Prolong Gold antifade reagent with DAPI | Thermo Fischer | #P36935 |
| iQ SYBR Green Supermix | Bio-Rad | #1708880 |
| **Critical Commercial Assays** |  |  |
| RNeasy Mini Kit | Qiagen | #74104 |
| iScript cDNA synthesis kit | Bio-Rad | #1708891 |
| **Experimental Models: Cell Lines** |  |  |
| Human nasal airway epithelial cells (HNEC) | This paper | N/A |
| Hek293T – R-spondin-1 mFc cell line | Trevigen Cat# 3710-001- | N/A |
| **Oligonucleotides** |  |  |
| Primers used for this manuscript |  | Table S4 |
| **Equipment** |  |  |
| Zeiss LSM800 confocal microscopy | Zeiss | N/A |
| Leica SP8X confocal microscope | Leica | N/A |
| Leica THUNDER imager | Leica | N/A |
| Leica TCS SP8 STED 3X microscope | Leica | N/A |
| Ussing chamber system | Physiologic Instruments | N/A |
| Voltage clamp | World Precision Instruments | #DVC-1000 |
| PowerLab | AD Instruments | #8/30 |
| Epithelial voltohmmeter EVOM2 | World Precision Instruments | N/A |
| Nanodrop spectrophotometer | Thermo Fisher | N/A |
| CFX96 real-time detection machine | Bio-Rad | N/A |
| **Software and Algorithms** |  |  |
| Zen Blue Software | Zeiss | https://www.zeiss.com/microscopy/int/products/microscope-software/zen.html |
| Prism 8 | GraphPad Software Inc. | https://www.graphpad.com/scientific-software/prism/ |
| Microsoft Excel | Microsoft Corporation | https://office.microsoft.com/excel |
| Adobe Illustrator | Adobe | https://www.adobe.com/nl/products/illustrator.html |
| LabChart 6 | AD Instruments | https://labchart.software.informer.com/6.0/ |
| CFX Manager 3.1 | BioRad | https://www.bio-rad.com/en-us/sku/1845000-cfx-manager-software?ID=1845000 |
| LAS X software | Leica | https://www.leica-microsystems.com/products/microscope- |
|  |  | software/p/leica-las-x-ls/ |
| ImageJ/FIJI |  | https://imagej.net/Fiji/Downloads |
| **Others** |  |  |
| Cytological brush | CooperSurgical | #C0004 |
| 12 mm<br>Transwell® with<br>0.4 µm Pore<br>Polyester<br>Membrane Insert | Corning | #3460 |
| 6.5 mm<br>Transwell® with<br>0.4 µm Pore<br>Polyester<br>Membrane Insert | Corning | #3470 |
| Matrigel | Corning | #354230 |
| BC isolation and<br>expansion<br>medium |  | Table S1 |
| Air-liquid<br>interface<br>differentiation<br>medium |  | Table S2 |
| Airway organoid<br>culture medium |  | Table S3 |

## Methods and Protocols

### Patient materials and sample collection

Nasal brushings were collected from healthy volunteers without respiratory tract symptoms (n=22 independent donors), and subjects with CF (n=22 independent donors) by a trained research nurse or physician, essentially as previously described ^33^. Sampling of adults was conducted using a cytological brush and without anesthetics. Nasal brushings of infants (<6 years old) were collected with a modified interdental brush ^34^. Samples were taken from the inferior turbinates of both left and right nostrils and stored in collection medium, consisting of advanced (ad)DMEM/F12, with GlutaMAX (1% v/v), HEPES (10 mM), Penicillin-Streptomycin (1% v/v), and Primocin (100 μg/mL). Nasal brushings were collected and stored with informed consent of all participants and was approved by a specific ethical board for the use of biobanked materials TcBIO (Toetsingscommissie Biobanks), an institutional Medical Research Ethics Committee of the University Medical Center Utrecht (protocol ID: 16/586). Nasal samples from infants with CF were collected as part of the Precision study (protocol ID: NL54885.041.16), which was approved by the Medical Research Ethics Committee of the University Medical Center Utrecht (Utrecht, The Netherlands). Resected inferior nasal turbinate tissue was obtained from a subject who underwent corrective surgery for turbinate hypertrophy at the Academic Medical Center in Amsterdam, the Netherlands. The tissue was accessible for research within the framework of patient care, in accordance with the “Human Tissue and Medical Research: Code of conduct for responsible use” (2011) (www.federa.org), describing the no-objection system for coded anonymous further use of such tissue without necessary written or verbal consent.

### Isolation and expansion of human nasal airway epithelial cells as 2D-cultures

Nasal cells were dissociated from the brush in the collection medium by scraping through a sterile P1000 pipette tip with the top cut off. After centrifugation (400 g for 5 min), the pellet was treated with TrypLE express enzyme, supplemented with 1x Sputolysin. The cells were incubated for 10 min at 37 °C and strained using a 100 µm strainer. After centrifugation (400 g for 5 min), the remaining pellet was used for isolation of human nasal epithelial cells (HNEC). HNEC were isolated and expanded on 6-well culture plates coated with collagen IV (50 µg/mL) and using BC isolation and expansion medium (Table S1) respectively. BC isolation medium contained additional antibiotics to suppress microbial outgrowth and was used during the first week of epithelial cell isolation, before switching to BC expansion medium. BC expansion medium included the gamma-secretase inhibitor DAPT, which reduces outgrowth of squamous cells at late passages. Growth factors (FGF7, FGF10, EGF, and HGF) were added freshly to the culture medium. Cultures were refreshed three times a week (Monday, Wednesday, Friday). After reaching 80-90% confluency, cells were passaged with TrypLE express enzyme and further expanded for one passage before freezing with CryoStor® CS10, supplemented with Y-27632 (5 µM) to create a master cell bank. For further use, a work cell bank was created of passage 2 cells, which was cryo-stored.

### Differentiation of 2D ALI-HNEC cultures

Expanded HNEC (passage =4-6) were cultured on transwell inserts (0.4 μm pore size polyester membrane), which were coated with PureCol (30 μg/mL). Cells were seeded in a density of 0.2 or 0.5*10^6^ cells on 24 or 12 wells inserts respectively and cultured in submerged conditions in BC expansion medium until reaching confluency. Afterwards, culture medium was changed with air-liquid interface (ALI)-differentiation medium (Table S2) supplemented with A83-01 (500 nM), and cells were additionally cultured in submerged condition for 1-2 days. Subsequently, culture medium at the apical side was removed and cells were further differentiated as ALI-cultures. After 3-4 days, cells were refreshed with ALI-diff medium without additional A83-01 and differentiated for at least 14 additional days at ALI-conditions. Medium was refreshed twice a week (Monday and Thursday or Tuesday and Friday), and the apical side of the cultures was washed with PBS once a week.

### Conversion of 2D differentiated ALI cultures into airway organoids

Differentiated ALI-cultures were washed at the apical surface with PBS and subsequently treated at the basolateral side with collagenase type II (1 mg/mL) diluted in adDMEM/F12. Cultures were incubated at 37 °C and 5% CO_2_ for 45-60 min until the epithelium detaches from the transwell insert. Next, the dissociated epithelial layer was transferred to a 15 ml tube in 1 ml adDMEM/F12 + 10 % (v/v) Fetal bovine serum (FBS), mechanically disrupted into smaller fragment by pipetting, and strained with a 100 µm filter. After centrifugation (at 400 g, 5 min), the epithelial pellet was resuspended in ice-cold 75% growth factor reduced Matrigel (v/v in airway organoid (AO) medium Table S3). Next, epithelial fragments were embedded in 30 µl Matrigel droplets on pre-warmed 24-well suspension plates. Droplets were solidified at 37 °C and 5% CO_2_ for 20-30 min, before adding 0.5 ml AO medium (Table S3). In optimized conditions for measuring CFTR modulator responses, AO culture medium was further supplemented with neuregulin-1β (NR, 0.5 nM) and interleukin-1β (IL-1β; 10 ng/ml). Besides, NR/IL-1β we furthermore examined the effects of culturing with other interleukins i.e. IL-13, IL-4, IL-10, and the growth factors: fibroblast growth factor 2, 7, 10, hepatocyte growth factor, and insulin-like growth factor 1, which did not improve the detection of CFTR modulator responses. AO medium was refreshed twice a week (Monday and Thursday or Tuesday and Friday).

### Forskolin-induced swelling (FIS) assay

ALI-derived NAOs were used in forskolin-induced swelling (FIS) assays, essentially as previously described with minor adaptation ^11,35^. In short, organoids were transferred 3-5 days after conversion in 96-well plates in 4 µl droplets of 75% matrigel (v/v in AO medium), containing approx. 25-50 structures. After solidification of droplets, 100 µl AO medium was added to each well. In optimized conditions for measuring CFTR modulator responses, AO culture medium was supplemented with neuregulin-1β (0.5 nM) and interleukin-1β (10 ng/ml). organoid swelling was conducted in quadruplicates. In indicated experiments, NAOs were pre-treated with CFTRinh-172 and GlyH101 (CFTRi, both 50 µM) or vehicle as negative control, for 4 h. CFTR correctors: VX-809, VX-661 (both 10 µM), VX-445 (5 µM), C1-18 (all 10 µM) or vehicle were pre-treated for 48 h. Before assessment of FIS, NAOs were stained with Calcein green AM (3 µM) for 30 min. Afterwards, organoids were stimulated with forskolin with indicated concentration. In cultures from subjects with CF, the CFTR potentiator VX-770 (10 µM) or vehicle was added together with forskolin. Swelling of NAOs was quantitated by measuring the increase of the total area of calcein green AM-stained organoids in a well during 15-min time intervals for a period of 2 h. Images were acquired with a Zeiss LSM800 confocal microscopy, using a 2.5 or 5x objective and experiments were conducted at 37°C and 95% O_2_/5% CO_2_ to maintain a pH of 7.4. Data was analysed using Zen Blue Software and Prism 8.

### Ussing chamber experiments

For open circuit ussing chamber measurements, transwell inserts (⍰ 12 mm) were mounted in the chamber device and continuously perfused at the apical and basal side with a Ringer solution of the following composition (mmol/L) 145 NaCl, 1.6 K_2_HPO_4_, 1 MgCl_2_, 0.4 KH_2_PO_4_, 1.3 Ca^2+^ Gluconate and 5 Glucose, and pH adjusted to 7.4. Following a 20 min stabilization period amiloride (20μM) was added to the apical side to block epithelial Na+ channel-mediated currents, followed by forskolin/IBMX (2μM/100µM), VX-770 (3μM) and CFTRInh-172 (30μM) were added sequentially. Transepithelial Voltage (V_te_) values were recorded at all times with Power Lab software (AD Instruments Inc.). Values for V_te_ were referred to the basal side of the epithelium and transepithelial resistance (R_te_) was determined by applying short intermittent pulses (0.5 μA/s), measuring pulsated deviations in V_te_ and accounting for the area of the inserts. An empty insert was previously recorded to correct the measured values. Short-circuit currents (I_eq-sc_) were calculated according to Ohm’s law from V_te_ and R_te_ (I_eq-sc_ = V_te_ / R_te_).

### RNA extraction, cDNA synthesis, and quantitative real time PCR

Total RNA was extracted from ALI-cultures using the RNeasy Mini Kit according to the manufacturer’s protocol. RNA yield was determined by a Nanodrop spectrophotometer and subsequently cDNA was synthesized by use of the iScript cDNA synthesis kit according to the manufacturer’s protocol. Quantitative real-time PCR (qPCR) was performed with specific primers (Table S4) using the iQ SYBR Green Supermix and a CFX96 real-time detection machine. CFX Manager 3.1 software was used to calculate relative gene expression normalized to the housekeeping genes *ATP5B* and *RPL13A* according to the standard curve method.

### Immunofluorescence staining and microscopy

2D expanded HNEC and ALI-HNEC cultures were fixed in 4% paraformaldehyde for 15 min, permeabilized in 0.25% (v/v) Triton-X in PBS for 30 min and treated with blocking buffer, consisting of 1% (w/v) BSA and 0.25% (v/v) Triton-X in PBS, for 60 min. Next, primary antibodies (1:500) in blocking buffer were added at the apical side and incubated for 1-2 h or overnight. Afterwards, cells were washed three times with PBS and incubated with secondary antibodies and Phalloidin (1:500) in blocking buffer for 30 min in dark, followed by three washings in PBS. Transwell membranes were subsequently cut from the inserts and placed on slides. All samples were mounted with Prolong Gold antifade reagent with or without DAPI. Resected nasal tissue and ALI-HNEC cultures were fixed with 4 % paraformaldehyde and embedded in paraffin after dehydration. After deparaffinization followed by antigen retrieval using citrate buffer (pH = 6) for 20 min, the 5 µm sections were permeabilized in 0.25% (v/v) Triton-X in PBS for 15 min, then treated with blocking buffer, consisting of 5% (w/v) BSA and 0.025% (v/v) Triton-X in PBS, for 30 min. Primary antibodies in blocking buffer, were incubated for 2 h, followed by incubation of secondary antibodies for 1 h. Samples were mounted with Prolong Gold reagent with DAPI. Organoids plated in 4 µl droplets of 75% matrigel (v/v) in a 96 wells plate, were fixed with 4 % paraformaldehyde for 10 min, and stained as previously described ^11,36^, using indicated primary antibodies. Images were acquired with a Leica SP8X confocal microscope, Leica THUNDER imager, and Leica TCS SP8 STED 3X microscope. Images were processed using LAS X software and ImageJ/FIJI.

### Quantification and statistical analysis

Swelling assays were conducted in quadruplicates for each experimental condition, and results are shown as mean ± SEM of independent subjects. Increases in the total surface area of all organoids in a single well is calculated as normalized swelling, relative to t=0, which is set as baseline of 100%. For swelling assays, statistical analysis was assessed with area under the curve (AUC) values (t=120 min). Analysis of differences was determined with a one/two-way repeated measurements ANOVA and Bonferroni *post-hoc* test or (un)paired Student’s t-test as indicated in the figure legends. Normal distribution was tested using the Shapiro Wilk test. Differences were considered significant at p< 0.05. Statistical analysis was conducted using Prism 8 (GraphPad Software Inc.).

## Acknowledgments

This work was supported by grants of the Dutch Cystic Fibrosis Foundation (NCFS, HIT-CF grant); Netherlands Organization for Health Research and Development (ZonMw); Health Holland (grant no 40-41200-98-9296); SRC 013 from CF Trust-UK; UIDB/04046/2020 and UIDP/04046/2020 centre grants (to BioISI), both from FCT/MCTES Portugal; and “HIT-CF” (H2020-SC1-2017-755021) from EU. This work is supported by the European Research Council (ERC Consolidator Grant 819219 to L.C.K.).

## Author contribution

Conception and/or design: GA, LR, CE, JMB; Acquisition, analysis, or interpretation of data: GA, LR, HR, EA, IS, WN, SV, BA, EK, JEB, SM, KW, HH, LK, MA, CE, JMB; Drafting the work or revising it critically for important intellectual content: GA, CD, MA, CE, JMB; Final approval of the version submitted for publication; GA, LR, HR, EA, IS, WN, SV, BA, EK, JEB, CD, SM, KW, HH, LK, MA, CE, JMB.

## Data availability

All data is provided with the manuscript.

