## Supplemental information for "Measuring cystic fibrosis drug responses in organoids derived from 2D differentiated nasal epithelia"

**Supplemental figure legends**

**Figure S1) Heterogeneity in tissue-derived airway organoid cultures.** (A) Brightfield images of airway organoids cultured according to Sachs et al.^1^, showing differences in organoid morphology, i.e. structures with lumen and solid colonies. (B) Immunofluorescence confocal imaging of airway organoids from the same culture, stained for the basal cell marker p63 (red) and ciliated cell marker β-tubulin IV (green). Image’s show (left) a solid colony consisting solely of p63^+^ basal cells, (middle) a pseudostratified structure with lumen but without ciliated cells, and (right) a large structure displaying ciliated cells. (C). Representative images (t=0 and 120 min after forskolin stimulation) of a FIS assay using HC airway organoids stained with calcein green-AM. Arrows indicate swelling of large-sized structures.

**Figure S2) Characterization of CFTR function in ALI-HNEC cultures.** Open circuit Ussing chamber tracings showing trans-epithelial voltage measurements (V_te_) from (A) HC and (B) CF F508del/F508del ALI-HNEC cultures. CF cultures were pre-treated with VX-809 or vehicle. During measurements, cells were treated sequentially with Amiloride, Fsk/IBMX , VX-770, and CFTRi. The bar corresponds to a 10 min frame.

**Figure S3) Characterization of nasal airway organoid swelling assay.**

(A and B) HC nasal airway organoids (NAOs) (n=2 independent donors) were pre-treated with bumetanide or vehicle for 4 h, followed by stimulation with forskolin (Fsk) and assessment of FIS. (C and D) Comparison between HC and CF organoid swelling (both n=6 independent donors) after stimulation with the calcium-activated chloride channel agonist E_act_. (E and F) CF F508del/F508del NAOs (n=8 independent donors) were pre-treated with VX-809 or vehicle control for 48 h. Subsequently, FIS was determined after acute stimulation with Fsk, (5 µM), together with VX-770 or vehicle control. Results are depicted as the percentage change in surface area relative to t=0 (normalized area) measured at 15-min time intervals for 2 h (means ± SD) (A, C,E) and as area under the curve (AUC) plots (t=120 min, means ± SD) ( B, D, F). Analysis of differences was determined with an unpaired (D) and paired t-test (F) . *** p<0.001.

**Figure S4) Optimization of CFTR modulator responses in NAOs with NR/IL-1β.** (A) NAOs from F508del homozygous subjects with CF (n=3 independent donors) were cultured without additional stimuli (Control), with neuregulin (NR), interleukin-1β (IL-1β), or combination of both (NR+IL-1β) for 5 days. At day 3, cultures were pre-treated with VX-809 or vehicle control for 48 h, and afterwards stimulated with forskolin (Fsk, 5 µM), in combination with VX-770 or vehicle control. Results are depicted as the percentage change in surface area relative to t=0 (normalized area) measured at 15-min time intervals for 2 h (means ± SD). (D) Brightfield images of CF F508del/F508del NAOs cultured in control conditions or with NR+IL-1β for 5 days. (E) Quantification of the mean organoid area (µm^2^, means ± SD) of control and NR+IL-1β cultured CF F508del/F508del NAOs (n=3 independent donors). (F, G) mRNA expression analysis of CF F508del/F508del NAOs (n=3-6 independent) that were left unstimulated (Control) or cultured with NR+IL-1β for 5 days. mRNA expression was determined of (F) *CFTR*, *ANO1*, *SLC26A9,* (all chloride channels), (G) *SPDEF* (secretory cells) and *FOXJ1* (ciliated cells). Results represent target mRNA expression normalized for the geometric mean expression of the housekeeping genes *ATP5B* and *RPL13A* (means ± SD). (H) Immunofluorescence staining of CF organoids (Control and cultured with NR+IL-1β for 5 days) was conducted of β-tubulin IV (ciliated cell) and MUC5AC (goblet cell) (both green). DAPI (blue) was used for nuclear staining. Phalloidin (red) was used for actin cytoskeleton staining. Analysis of differences was determined with an paired t-test (C,E,F,G). * p<0.05, ** p<0.01.

**Figure S5) Validation of CFTR modulator responses in CF NAOs cultured with NR/IL-1β.** (A) Brightfield images (top) and images of calcein green AM-stained (bottom) NAOs cultured ith NR/IL-1β, showing FIS in HC and CF F508del/F508del cultures. FIS of CF NAOs was determined in combination with vehicle or VX809/VX-770. Images were taken at t=0, and 120 min. (B) VX-809/VX-770 modulator response measurement in NAOs of individual donor with a CF F508del/F508del genotype (means ± SD, measurements in quadruplicates). (C) CF F508del/F508del NAOs (n=1 donor, means ± SD, measurements in quadruplicates) were pre-treated with VX-809, or the CFTR correctors C1-18 for 48 hours. Afterwards, FIS was measured after stimulation with forskolin (Fsk) and VX-770. Vehicle was used as control. (D) Images showing FIS of NR+IL-1β cultured NAOs from individuals with CF and F508del/S1251N genotype. NAOs were stimulated with Fsk and vehicle (top panels) or VX-770 (bottom panels). (E) Images of FIS conducted in calcein green AM-stained CF F508del/F508del NAOs cultured with NR/IL-1β, pre-treated with vehicle (top panels) or triple CFTR modulator combination VX-661/VX-445/VX-770 (bottom panels). (D and E) Images were taken at t=0, and 120 min. (F and G) CF F508del homozygous NAOs (n=3 independent donors) were pre-treated with vehicle, VX-445, or a combination of VX-661 and VX-445 for 48 h. Next, cultures were pre-treated with CFTR inhibitors CFTRinh-172 and GlyH101 or vehicle for 4 h, followed by stimulation with Fsk together with VX-770 or vehicle as indicated. Results are depicted as percentage increase in normalized area in time (means ± SD) (F) and AUC plots (t=120 min, means ±SD) (B, C, F, G). Analysis of differences was conducted with a paired t-test (E, G) * p<0.05.

**Supplemental video S1: Live imaging of an ALI-cultured derived nasal airway organoids.** Video recording of a CF nasal airway organoid showing beating cilia at the luminal side of the structure and accumulated mucus.
