## Supplementary figures and images for "Measuring cystic fibrosis drug responses in organoids derived from 2D differentiated nasal epithelia"

### Figure S1

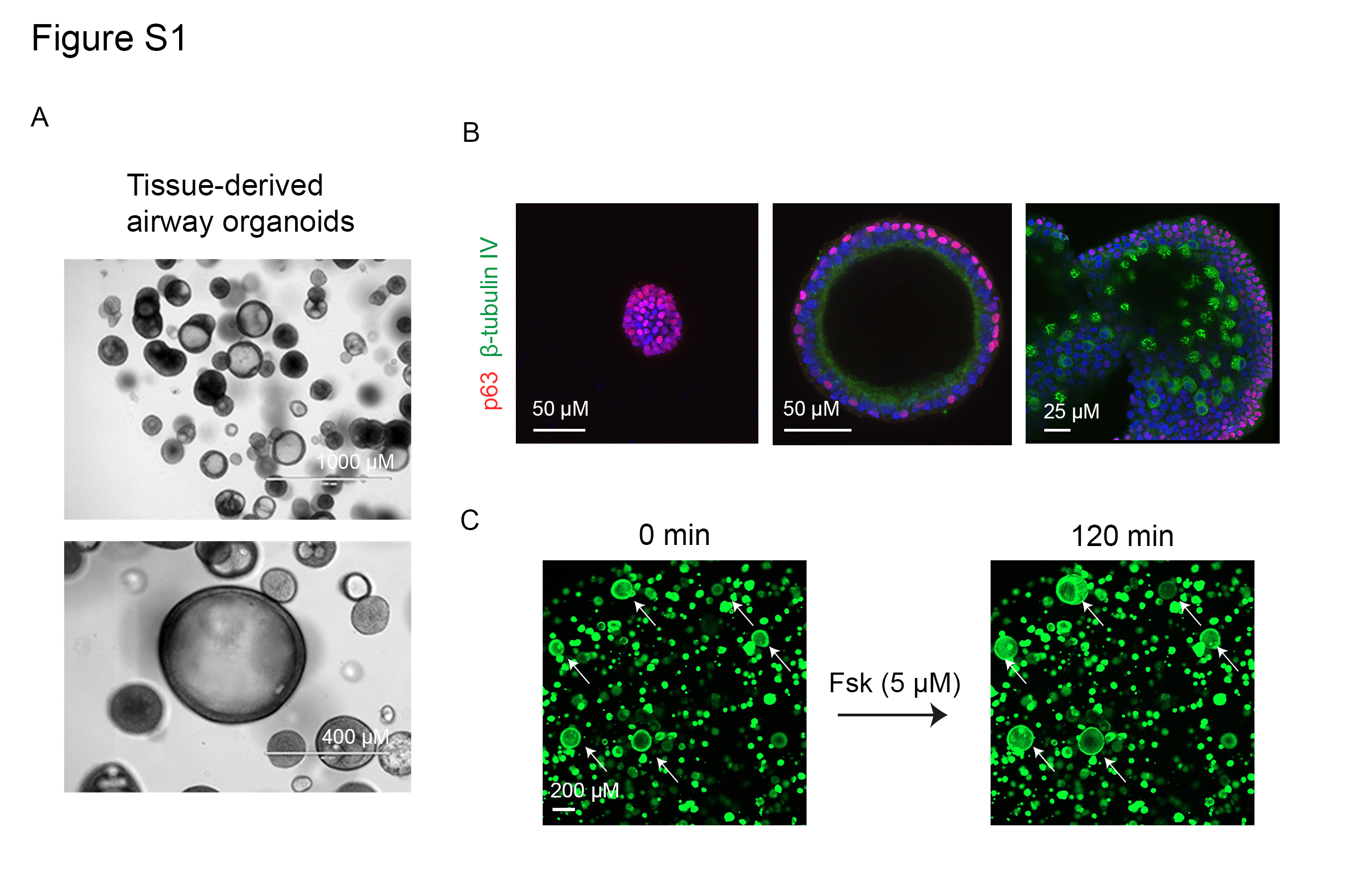

### Figure S2

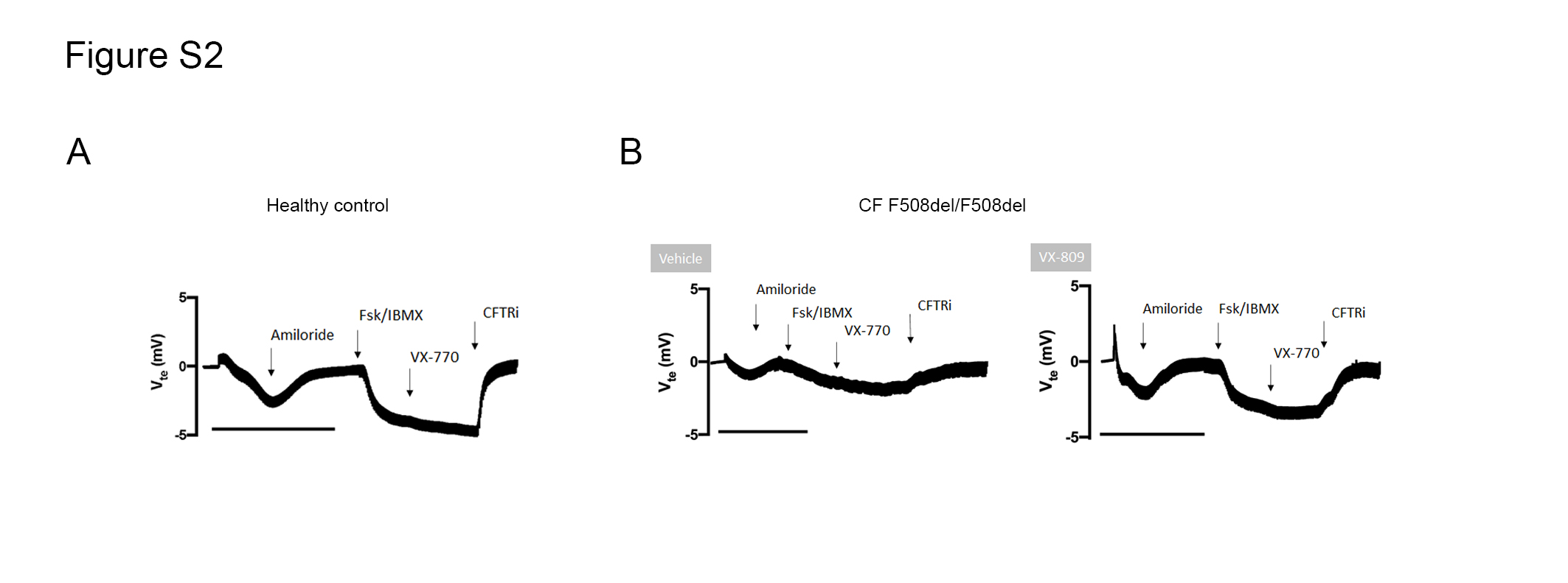

### Figure S3

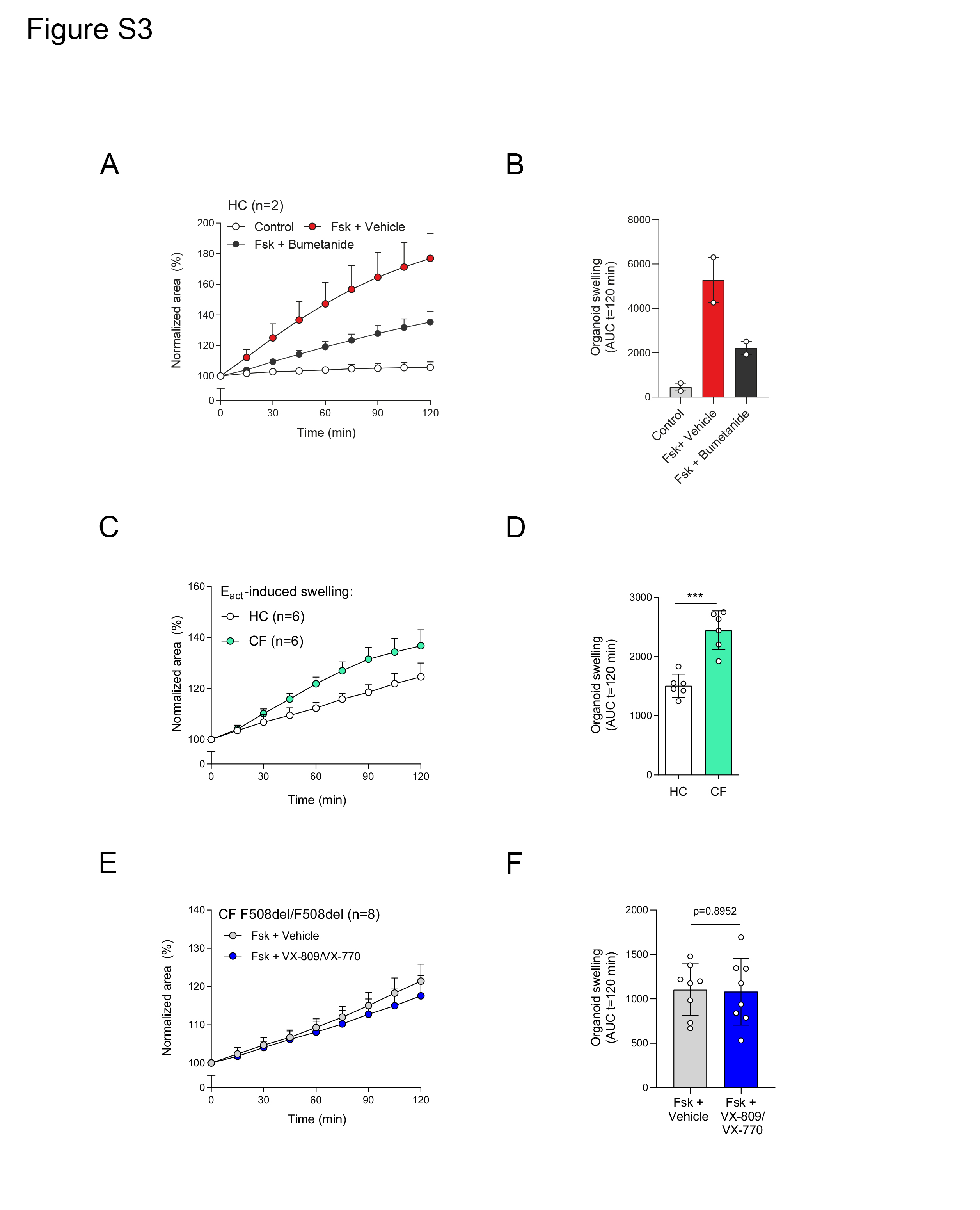

### Figure S4

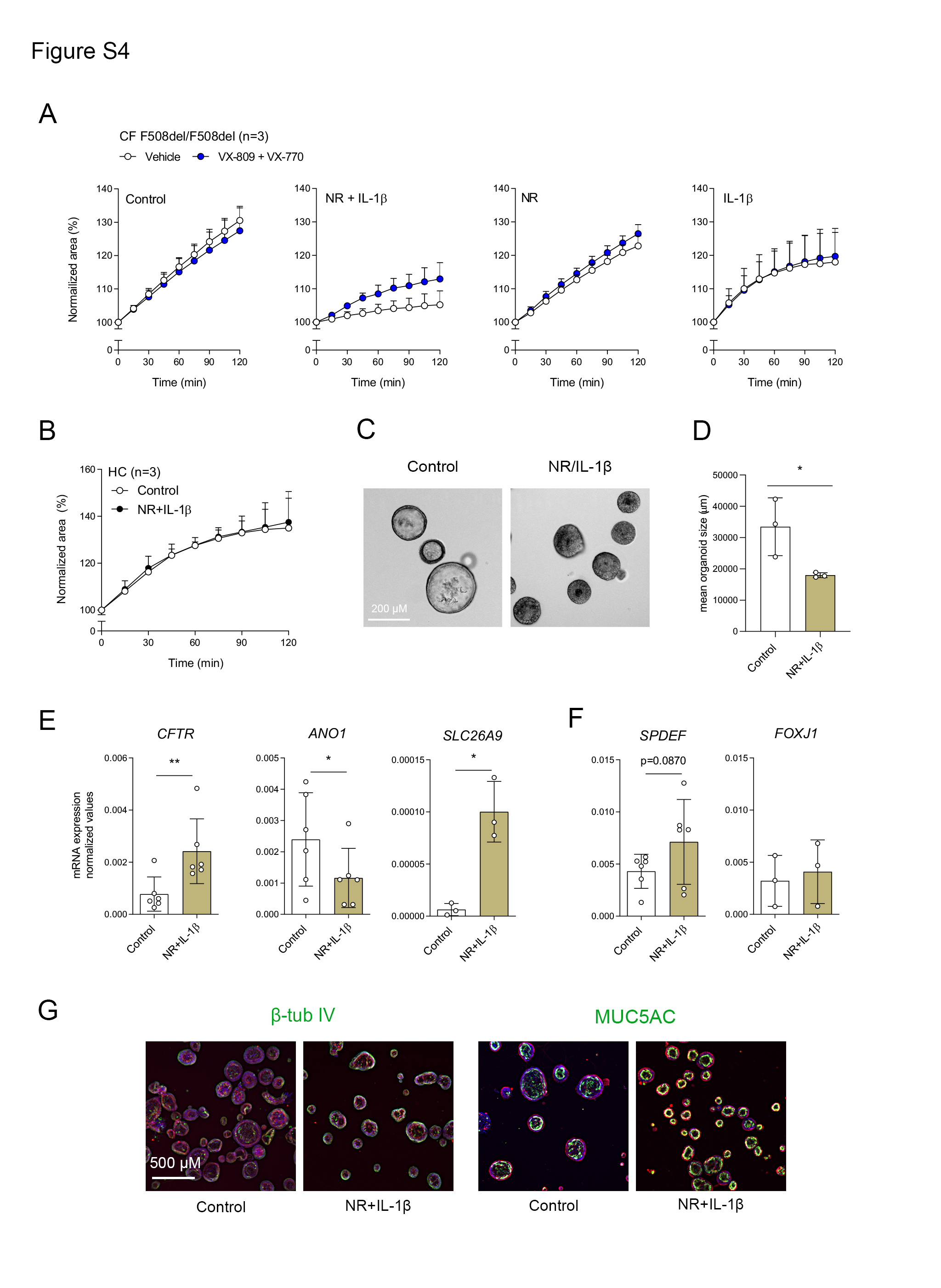

### Figure S5

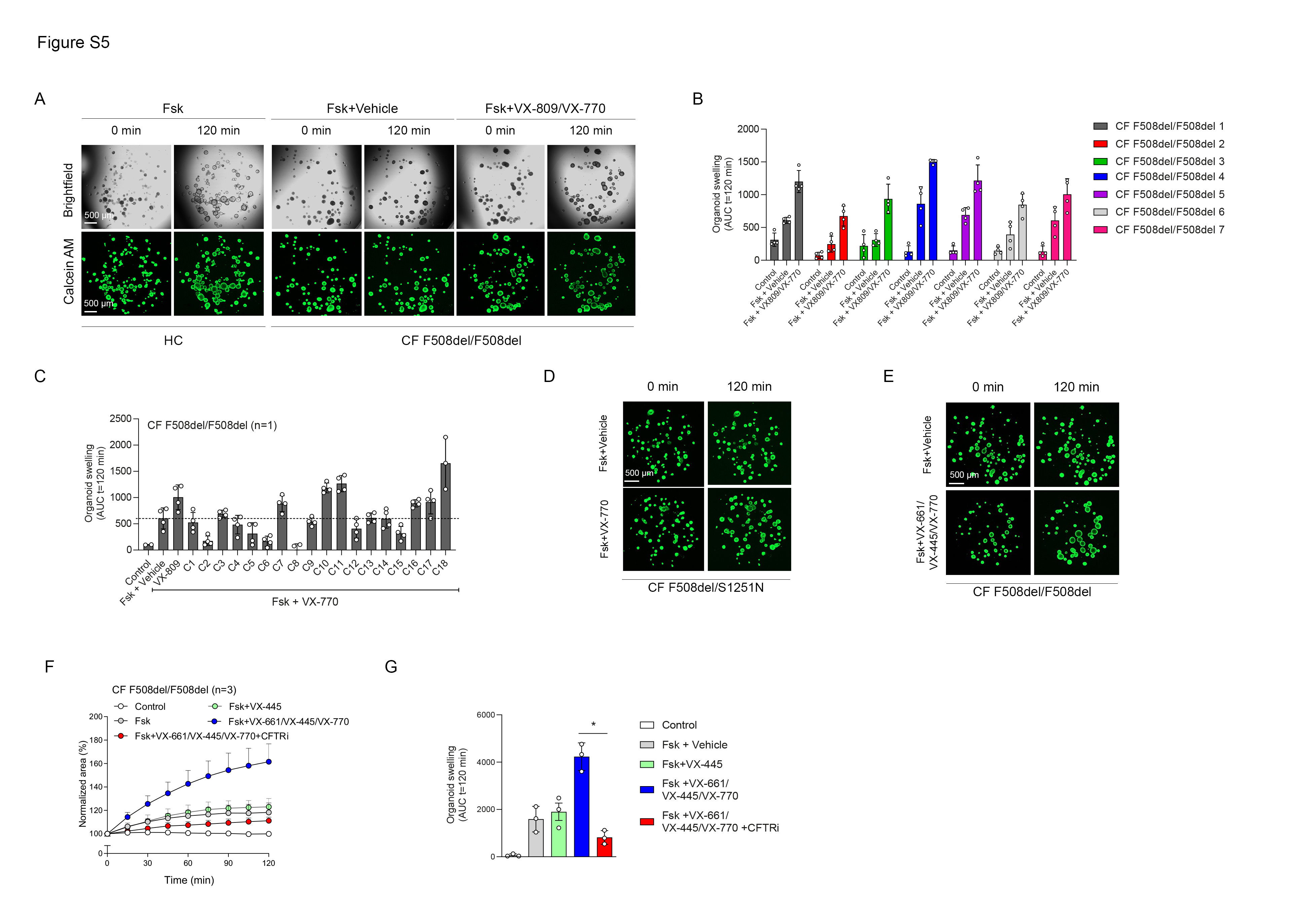
